# Dasabuvir inhibits human norovirus infection in human intestinal enteroids

**DOI:** 10.1101/2021.07.02.450857

**Authors:** Tsuyoshi Hayashi, Kosuke Murakami, Junki Hirano, Yoshiki Fujii, Yoko Yamaoka, Hirofumi Ohashi, Koichi Watashi, Mary K. Estes, Masamichi Muramatsu

## Abstract

Human noroviruses (HuNoVs) are acute viral gastroenteritis pathogens that affect all age groups, yet no approved vaccines and drugs to treat HuNoV infection are available. In this study, with a human intestinal enteroid (HIE) culture system where HuNoVs are able to replicate reproducibly, we screened an antiviral compound library to identify compound(s) showing anti-HuNoV activity. Dasabuvir, which has been developed as an anti-hepatitis C virus agent, was found to inhibit HuNoV infection in HIEs at micromolar concentrations. Dasabuvir also inhibited severe acute respiratory syndrome coronavirus 2 (SARS-CoV-2) and human A rotavirus (RVA) infection in HIEs. To our knowledge, this is the first study to screen an antiviral compound library for HuNoV using HIEs and we successfully identified dasabuvir as a novel anti-HuNoV inhibitor that warrants further investigation.

## Main text

Human noroviruses (HuNoVs) cause acute gastroenteritis and foodborne diseases among all age groups worldwide. HuNoVs often cause an economic burden to societies due to health care costs and loss of productivity, and therefore pose a public health concern. Noroviruses are non-enveloped viruses possessing a positive sense, single stranded RNA genome whose length is approximately 7.5kb long. They are genetically classified in 10 genogroups (GI-GX) and further divided into 48 genotypes based on their capsid and polymerase gene sequences (Chhabra et al., 2019). Among those, GII.4 genotype is the most frequently distributed and causes outbreak in humans globally (Cannon et al., 2021; Mallory et al., 2019).

Since a robust culture system to allow HuNoV replication was not established for almost 50 years, there is no established treatment options such as vaccines or antiviral regimens available. Recently, several HuNoV successive cultivation models employing a human B cell line (Jones et al., 2014), tissue stem cell-derived human intestinal enteroids (HIEs) (Ettayebi et al., 2016), human induced pluripotent stem cell-derived intestinal organoids (Sato et al., 2019), and zebrafish larvae (Van Dycke et al., 2019) have been developed. The stem cell-derived HIE system is currently used by researchers worldwide to study HuNoV biology and inactivation strategies (Alvarado et al., 2018; Chan et al., 2021; Costantini et al., 2018; Ettayebi et al., 2021; Hosmillo et al., 2020; Lin et al., 2020; Murakami et al., 2020), although, to our knowledge, compound screens for identifying HuNoV antiviral agents have not been reported.

Drug repurposing is a time-saving, affordable strategy to discover new therapeutic uses for approved or developing drugs to treat other disease(s) apart from their original use (Low et al., 2020). This strategy is being widely utilized to establish effective therapeutics for treatment of coronavirus disease 2019 (COVID-19). Indeed, numerous antiviral drugs including remdesivir, ivermectin, or nelfinavir has been identified as promising candidates for SARS-CoV-2 (Low et al., 2020; Watashi, 2021). Here, with the HIE culture system, we screened an antiviral compound library composing 326 bioactive substances, including those targeting influenza virus, human immunodeficiency virus (HIV), or Hepatitis C virus (HCV) to reassess their effect on HuNoV infection.

Three dimensional (3D) HIEs were dissociated, and plated on collagen-coated 96 well plates to prepare two-dimensional (2D) monolayers (**Fig. 1A**). The cells were then differentiated by culturing them in differentiation medium, which does not include Wnt3A and R-spondin to support HIE’s stemness. The differentiated HIE monolayers were inoculated with GII.4 HuNoV in the presence of each compound dissolved in DMSO for 1 hr at 37°C. DMSO was added to the wells without compound (DMSO control). For this screening step, one well was used to analyze each compound (n = 1). The cells were washed, and cultured in differentiation medium containing the compound for 24 hrs. The infected cells and supernatant were then harvested, and viral replication was evaluated by RT-qPCR analysis to determine the HuNoV RNA genome equivalents (GEs) (**Fig. 1B**). Cytotoxicity for each compound was also determined by LDH assay. First, to evaluate the reproducibility of our HIE system with respect to HuNoV growth throughout the screening, we plotted the level of viral GEs at 1 or 24 hr post-infection (hpi) from 7 independent experiments that were used for the compound screen. The fold changes of viral GEs between 1 and 24 hpi in each experiment ranged from 65 to 230 (mean ± s.d.; 104.3 ± 58.3). A positive control, 2’-C-Methylcytidine (2-CMC) completely blocked viral infection without any cytotoxicity in all experiments, consistent with a previous report (**Fig. 1C and D**) (Hosmillo et al., 2020). These results demonstrate that our HIE cultivation system reproducibly supports HuNoV replication, and is suitable for evaluating the effect of the compounds against HuNoV.

**Figure 1.**
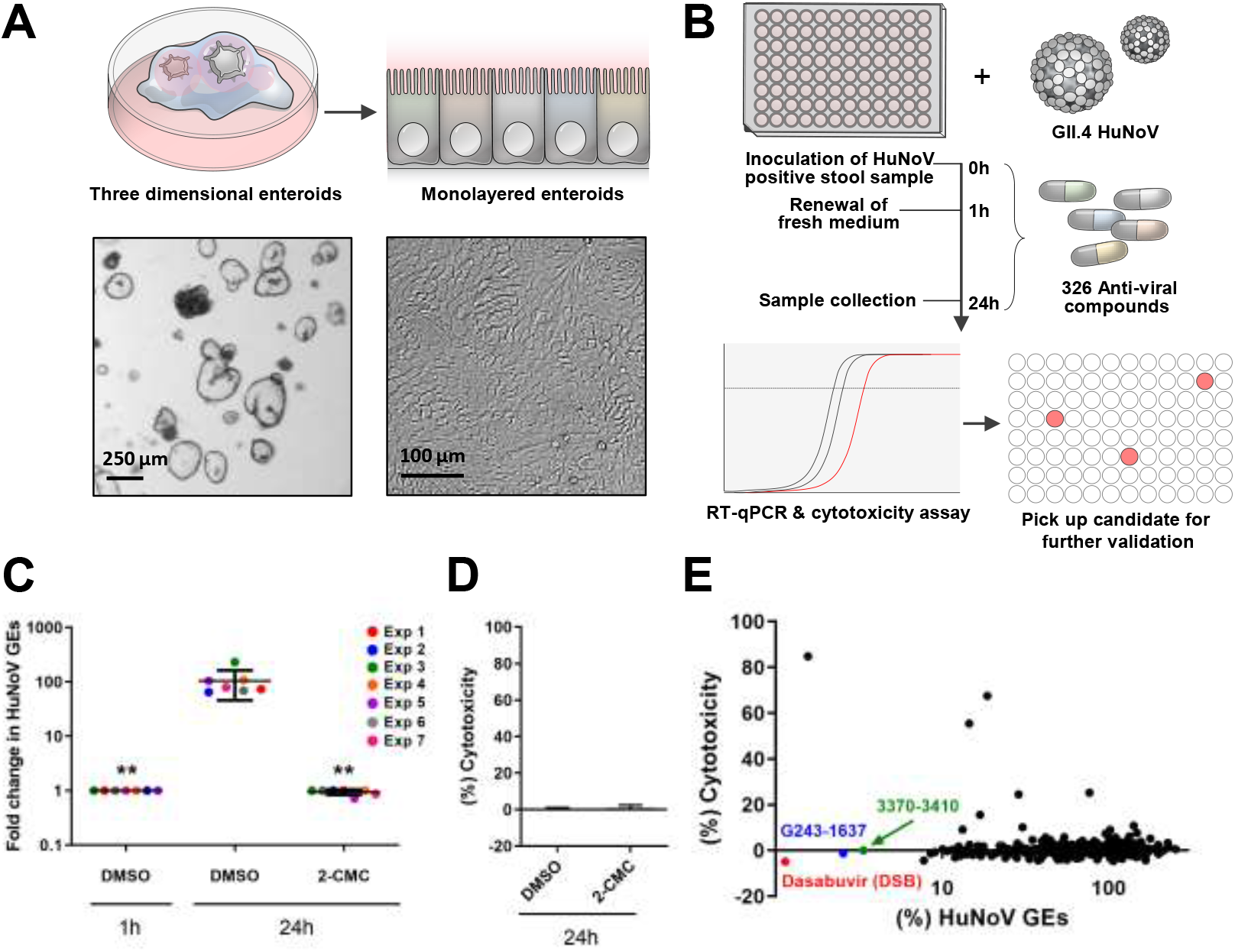
Screening for antiviral compounds that inhibit human norovirus infection in human intestinal enteroids. (A-B) Schematic illustrations of compound screening. Three dimensional HIEs (J2) were trypsinized to single cells, and plated in 96-well plate to culture them as 2D monolayers. Differentiated HIE monolayers were then inoculated with GII.4 HuNoV-containing stool filtrates in the presence of the compounds (10 μM, n=1). After 1 hr incubation at 37°C, the cells were washed, and cultured in differentiation medium containing the compounds (10 μM) in 100 μL volume until 24 hrs post infection (hpi). Viral RNA extracted from the cells and 75 μL of supernatant at 1 or 24 hpi was subjected to RT-qPCR to measure viral genome equivalents (GEs). The rest of the supernatant was subjected to LDH assay to measure cytotoxicity. (C) HuNoV replication in J2 monolayers throughout the screening. We performed 7 experiments to screen all 326 compounds. DMSO and 2’-C-Methylcytidine (2-CMC, 389 μM) were used as controls in every test. Viral GEs in DMSO and 2-CMC treated samples at 24 hpi were normalized to DMSO control at 1 hpi. ** *p* < 0.01 *versus* DMSO control at 24 hpi, one-way ANOVA followed by Dunnett’s multiple-comparison test. (D) Cytotoxicity of 2-CMC in J2 monolayers at 24 hpi. Results were normalized to DMSO control. (E) Scatter plot of the % HuNoV GEs *vs* % cytotoxicity for all tested compounds. Results were normalized to DMSO control. Dasabuvir (DSB), red; G243-1637, blue; 3370-3410, green spots.

We next determined the relative percentages of HuNoV GE and cytotoxicity at 24 hpi by normalizing the data of compound-treated cells to DMSO-treated cells. The screening results were plotted as (%) HuNoV GE *vs* (%) Cytotoxicity (**Fig. 1E**). We selected 3 compounds, Dasabuvir (DSB), G243-1637, and 3370-3410, which reduced HuNoV GEs by 95% without cytotoxicity for further validation (**Fig. 1E and Table S1**). For validation purpose, we repeated the experiment with technical 3 replicates and confirmed the reproducible inhibitory effect of DSB and G243-1637 against HuNoV infection (**Fig. S1**). We selected DSB which showed strongest inhibitory effect for further studies. DSB has been developed as one of direct-acting anti-HCV drugs which targets HCV NS5B RNA-dependent RNA polymerase (RdRp) (Trivella et al., 2015). So far, there have been no report regarding any antiviral effects on HuNoV.

Next, we performed additional experiments, with DSB at varying concentrations ranging from 3.125 μM to 50 μM to calculate EC_50_ and CC_50_ values. DSB treatment alone showed no cytotoxicity, except for the highest concentration (50 μM), which showed a 17% reduction of cellular ATPs (cell viability) or 10% increase of LDH release (cytotoxicity), as compared to the DMSO control (**Fig. S2**). Again, DSB did not induce cytotoxicity, except at the highest dose (50 μM) in J2 monolayers infected with GII.4 HuNoV, whereas it showed a dose-dependent inhibition of viral replication with an EC_50_ value of 11.71 μM (**Fig. 2A**). To ascertain the authenticity of DSB’s inhibitory effect, we repeated the experiment with the identical compound from different resources and found that the results were comparable (**Fig. S3**). Dose-dependent reduction of viral replication by DSB was also observed in J2 HIE monolayers infected with a different HuNoV strain GII.3 (**Fig. 2B)**. We further assessed DSB’s inhibitory effect using J3 HIEs established from an independent donor following the infection of GII.3 or GII.4 HuNoV and observed the same trends (**Fig. 2C and D)**. Taken together, DSB exerted an inhibitory effect on two HuNoV genotypes and HIEs established from distinct individuals, strongly suggesting that the effect is neither genotype- nor HIE (donor)-dependent.

**Figure 2.**
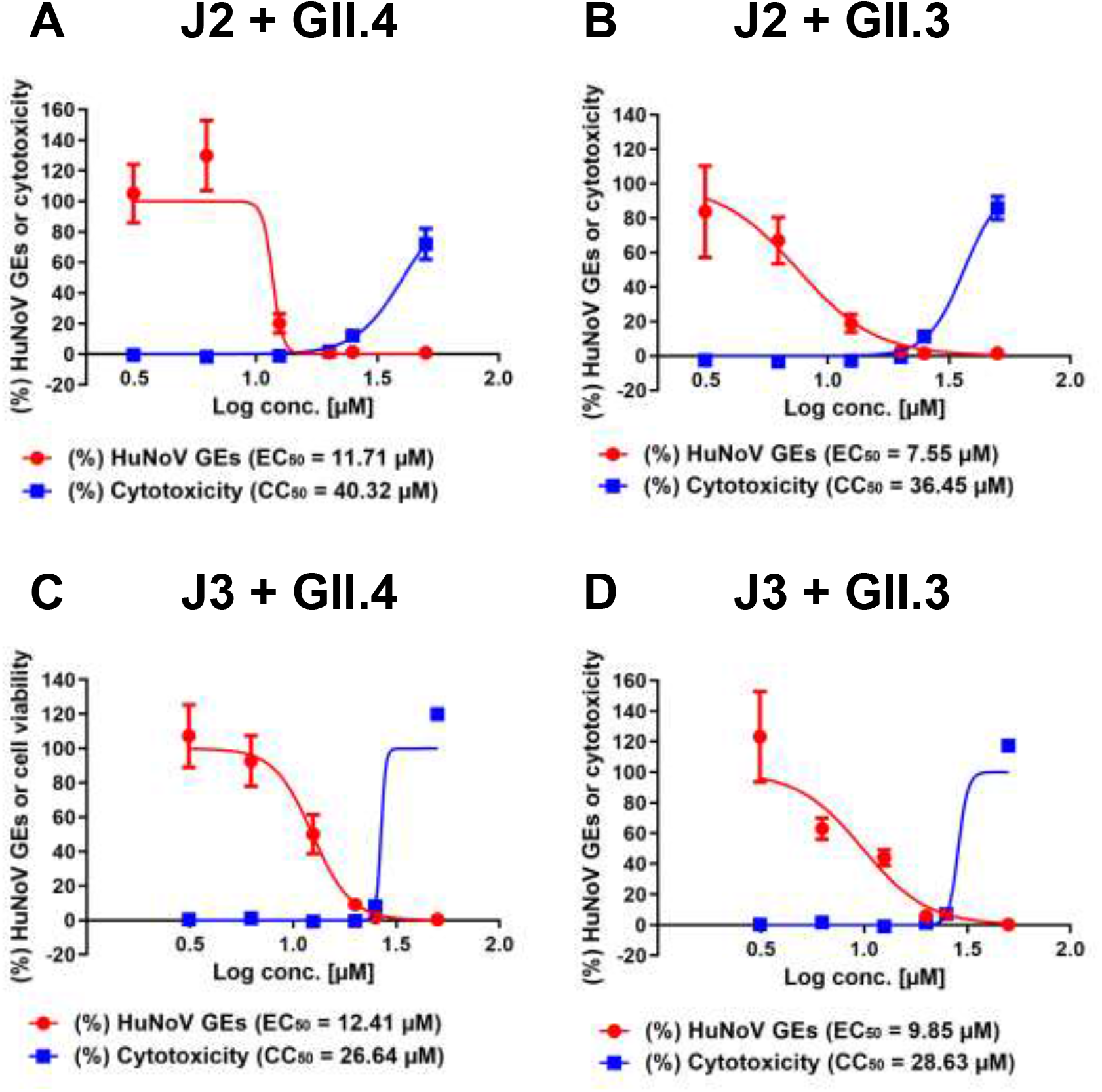
Effect of dasabuvir on HuNoV infection in HIE monolayers. J2 (A-B) or J3 (C-D) HIE monolayers were inoculated with GII.4 (A and C) or GII.3 (B and D) HuNoV-containing stool filtrate in the presence of DSB at the indicated concentrations, and were cultured until 20 hpi. The percentages of HuNoV GEs (red lines) and cytotoxicity (blue lines) were determined as in Fig. 1, and were normalized to the DMSO control. Values represent the mean ± s.d. (n ≥ 6). EC_50_; 50% effective concentration, CC_50_; 50% cytotoxic concentration.

Next, we tested the effect of DSB on the infection of human A rotavirus (RVA) and SARS-CoV-2, both of which has been previously reported to be able to infect and replicate HIEs (Lamers et al., 2020; Saxena et al., 2016; Zou et al., 2019). Two concentrations of DSB were used in these studies; non-effective (6.25 μM) and effective concentration (20 μM) against HuNoV, respectively. As shown in **Fig. 3A and B**, DSB showed an antiviral effect (2.75-fold decrease) on RVA infection at a concentration of 20 μM, while it almost completely inhibited (26.9-fold decrease) SARS-CoV-2 infection in J2 HIE monolayers. We also confirmed DSB’s inhibition with no cytotoxicity using VeroE6/TMPRSS2 cells which are highly susceptible to SARS-CoV-2 infection (**Figs. 3C, D, and S4**) (Matsuyama et al., 2020).

**Figure 3.**
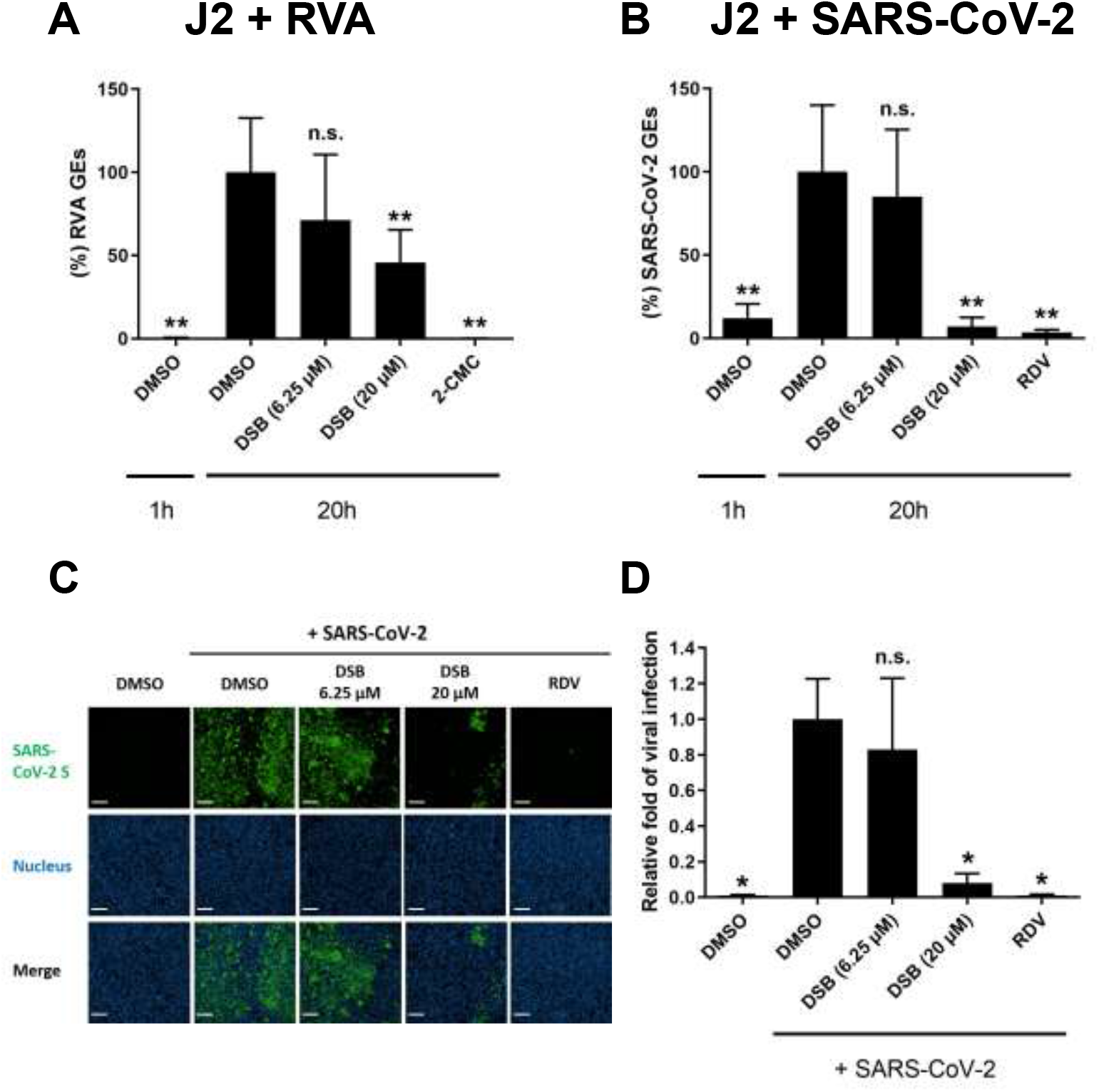
Dasabuvir inhibits SARS-CoV-2 infection in J2 HIE monolayers and VeroE6/TMPRSS2 cells. (A-B) J2 HIE monolayers were inoculated with RVA (A) and SARS-CoV-2 (B) in the presence of the indicated compounds, and were cultured until 20 hpi. 2-CMC (389 μM) or remdesivir (RDV, 10 μM) were used as positive controls. The percentage of viral GEs were determined by RT-qPCR, and were normalized to the DMSO control at 20 hpi. Values represent the mean ± s.d. (n ≥ 5). (C-D) VeroE6/TMPRSS2 cells were left-uninfected or infected with SARS-CoV-2 for 20 hrs in the presence of the indicated compounds. The cells were then stained with anti-SARS-CoV-2 Spike RBD monoclonal antibody and DAPI followed by imaging analysis. (C) Representative fluorescence images showing SARS-CoV-2 S protein (green) and cell nucleus (blue). Scale bar, 200 μm. (D) The percentages of infected cells were normalized to those of DMSO-treated cells infected with SARS-CoV-2 at 20 hpi. Values represent the mean ± s.d. (n ≥ 8). ** *p* < 0.01 *versus* DMSO control at 20 hpi, one-way ANOVA followed by Dunnett’s multiple-comparison test. n.s., not significant (*p* > 0.05).

The mechanism of action for the virus inhibitions by DSB remains to be elucidated. DSB is a non-nucleotide inhibitor of HCV NS5B RdRp that likely binds to palm domain of NS5B and thereby prevents elongation of the nascent viral genome (Kati et al., 2015). Therefore, it might also target the RdRp of other viruses such as HuNoV and SARS-CoV-2, possibly because of the presence of conserved sequences being targeted by DSB. Indeed, there is a report showing that DSB partially inhibits RdRp activity of Middle East respiratory syndrome coronavirus (MERS-CoV) (Min et al., 2020). Targeting viral protease might be another scenario for the inhibition; a very recent virtual screening study predicted that dasabuvir has a potential to inhibit 3-chymotrypsin-like protease (3CL^PRO^) of SARS-CoV-2 (Jade et al., 2021).

With a HCV subgenomic replicon system, DSB inhibits HCV of genotype 1 with EC_50_ values of < 10 nM (Kati et al., 2015). In contrast, DSB inhibits HuNoV infection with EC_50_s ranging between 7.55 and 12.41 μM (**Fig. 2**), which is comparable to its effectiveness to inhibit RdRp activity of MERS-CoV-2 (Min et al., 2020) or infection of vector-borne flaviviruses (Stefanik et al., 2020). This implies that higher concentration is required to exert an antiviral effect on non-HCV viruses, possibly due to lower binding efficiency of DSB to non-HCV RdRp(s) or unknown mechanism of inhibitory action.

In summary, through the screening of an antiviral compound library, we identified DSB as a novel HuNoV inhibitor that warrants further clinical investigation. To our knowledge, this was a first time to identify ‘*bona fide*’ anti-HuNoV agents using the HIE culture system. Our study also shed light on the usefulness of the HIE platform for investigating of anti-HuNoV agents and/or host factors regulating HuNoV infection, which will contribute to better understanding of HuNoV lifecycle and development of vaccine and antiviral regimens.

## Acknowledgements

This study was supported by grants from by the Japan Society for the Promotion of Science KAKENHI Grant JP20K07520 (to T.H.); the Japan Agency for Medical Research and Development (AMED) Grants JP21fk0108149 (to K.M. and T.H.), JP21fk0108102 (to M.M. and K.M.), JP21fk0108121 (to K.M.), JP21wm0225009 (to M.M.), JP21fk0108121 (to Y.F.), and NIH Grant PO1 AI057788 (to M.K.E.). We thank Drs. Hiroyuki Shimizu, Shutoku Matsuyama and Noriyo Nagata (National Institute of Infectious Diseases) for technical assistance.

---

**Figure S1.**
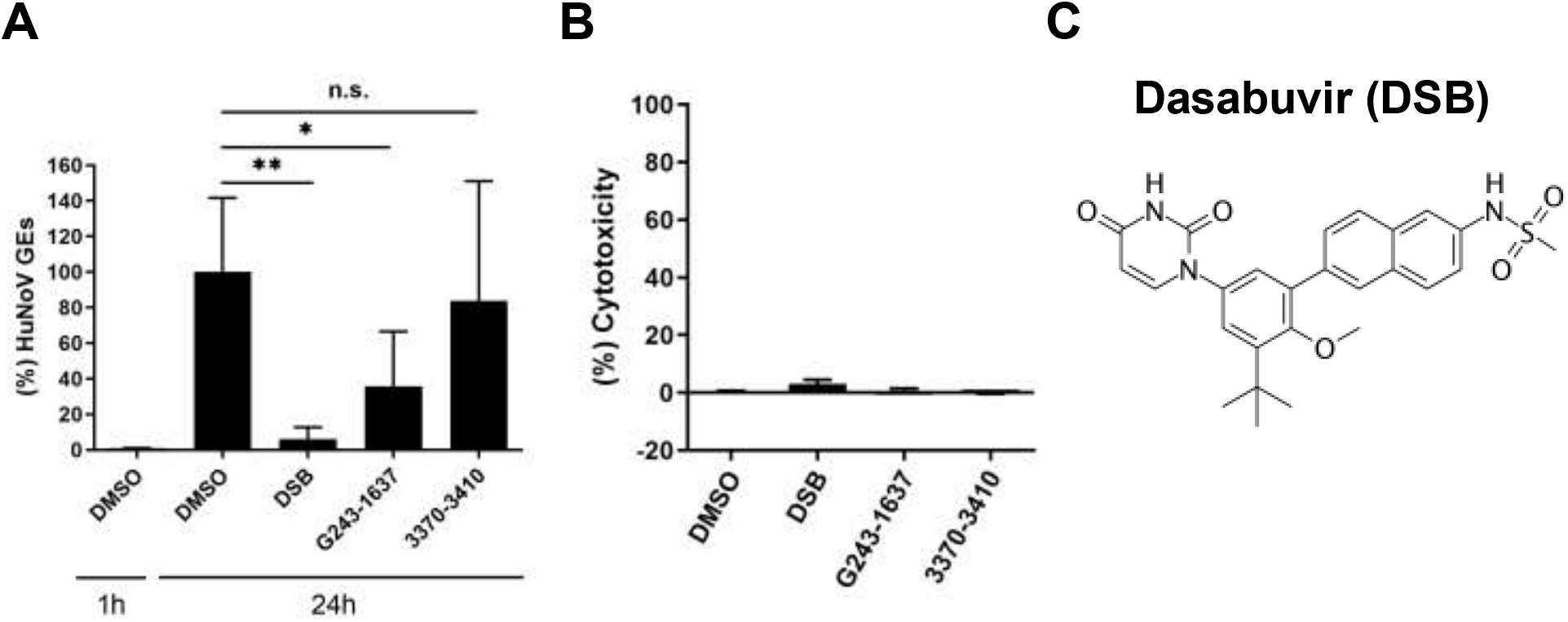
Validation of the selected 3 compounds in J2 HIE monolayers. J2 HIE monolayers were inoculated with GII.4 HuNoV-containing stool filtrate in the presence of the indicated compounds (10 μM), and were cultured until 24 hpi. The percentages of HuNoV GEs (A) and cytotoxicity (B) for each compound were determined, as in Fig. 1. Values represent the mean ± s.d. (n ≥ 3). ** *p* < 0.01, * *p* < 0.05 *versus* DMSO control at 24 hpi, two-tailed Student *t* test. n.s., not significant (p > 0.05). (C) Chemical structure of Dasabuvir.

**Figure S2.**
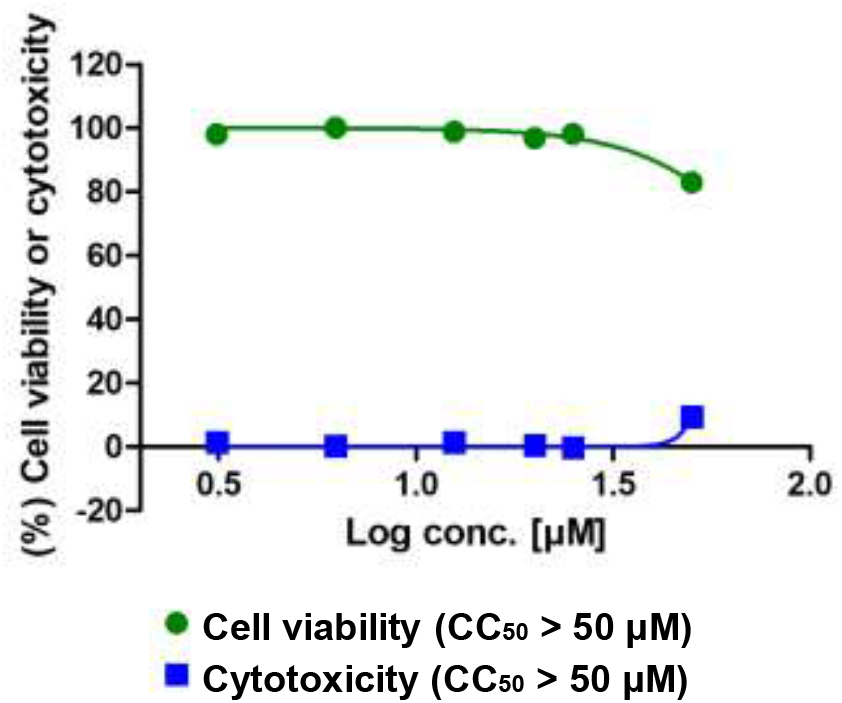
Cell viability and cytotoxicity of dasabuvir in J2 HIE monolayers. J2 HIE monolayers were treated with DSB at the indicated concentrations for 20 hr. Cell viability or cytotoxicity was measured using CellTiter-Glo^®^ Luminescent Cell Viability Assay or Cytotoxicity LDH Assay Kit-WST, respectively. Results were normalized to DMSO control. Values represent the mean ± s.d. (n = 7).

**Figure S3.**
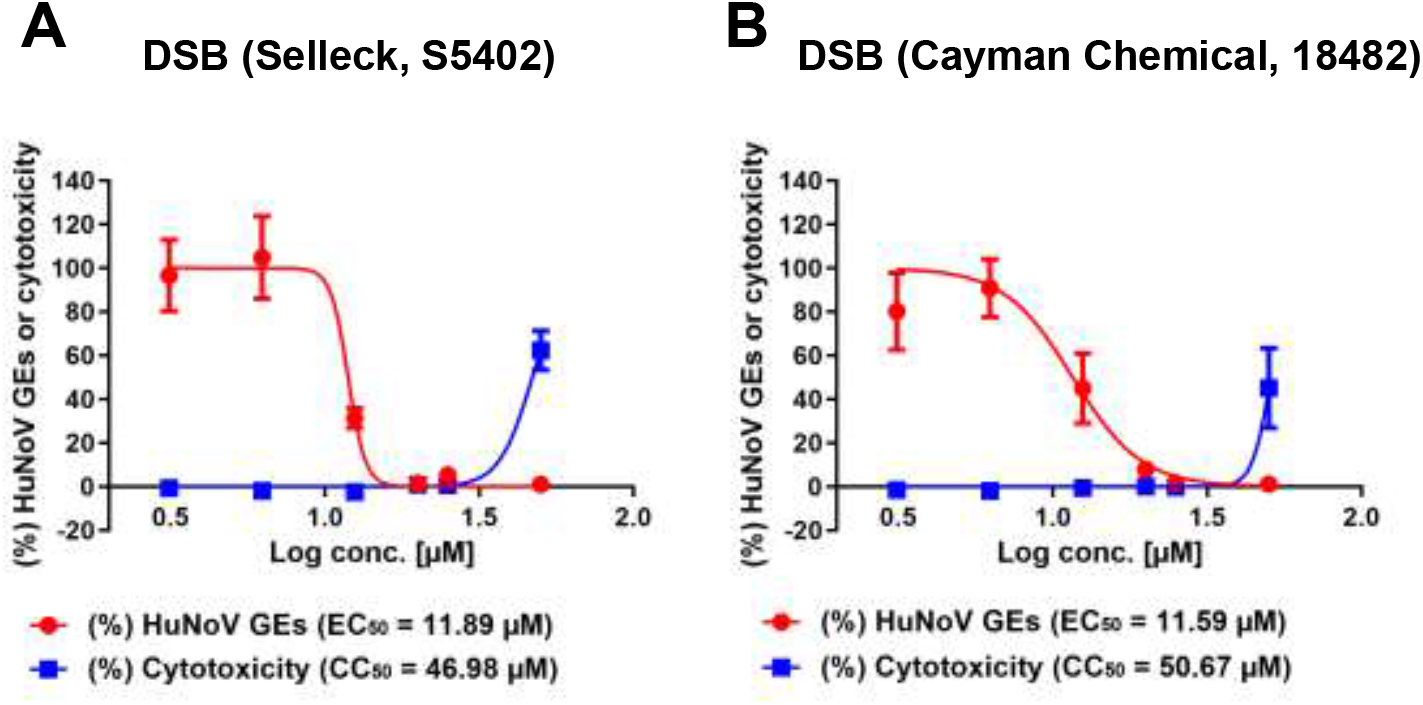
Inhibitory effect of dasabuvir purchased from different resources on HuNoV infection in J2 HIE monolayers. J2 HIE monolayers were inoculated with GII.4 HuNoV-containing stool filtrate in the presence of dasabuvir (DSB) purchased from different resources at the indicated concentrations, and cultured until 20 hpi. The percentages of HuNoV GEs and cytotoxicity were determined and were normalized to the DMSO control. Values represent the mean ± s.d. (n = 4).

**Figure S4.**
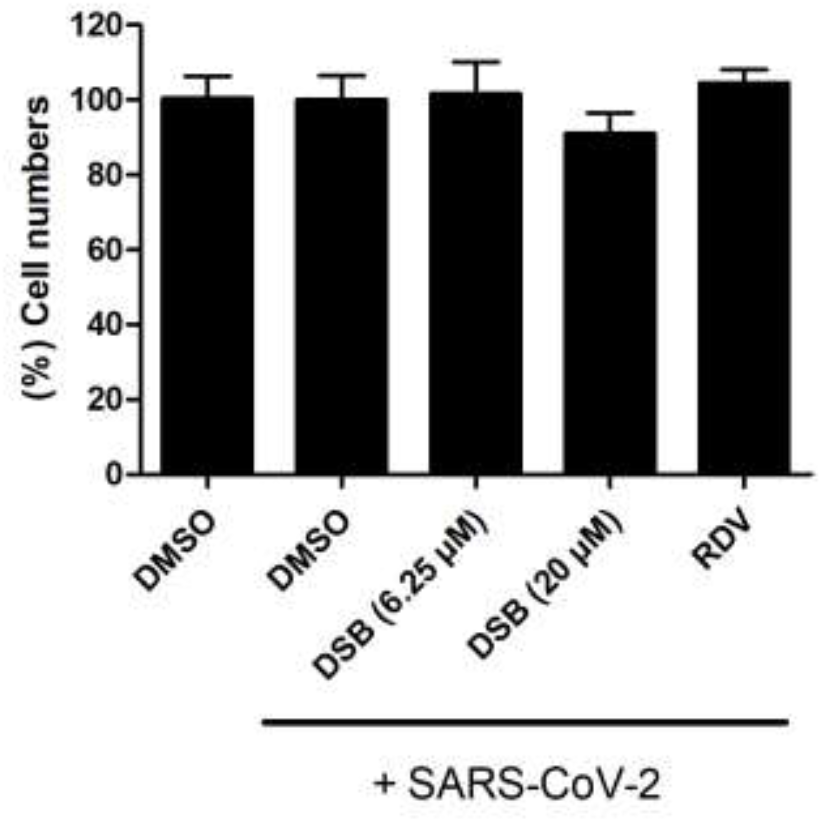
Cell viability of dasabuvir in VeroE6/TMPRSS2 cells. VeroE6/TMPRSS2 cells were left-uninfected or infected with SARS-CoV-2 for 20 hrs followed by immunofluorescence analysis, as described in Material and Methods. The cell numbers were determined by counting cell nucleus. Results were normalized to DMSO-treated cells infected with SARS-CoV-2. Values represent the mean ± s.d. (n = 8).

**Table S1.**
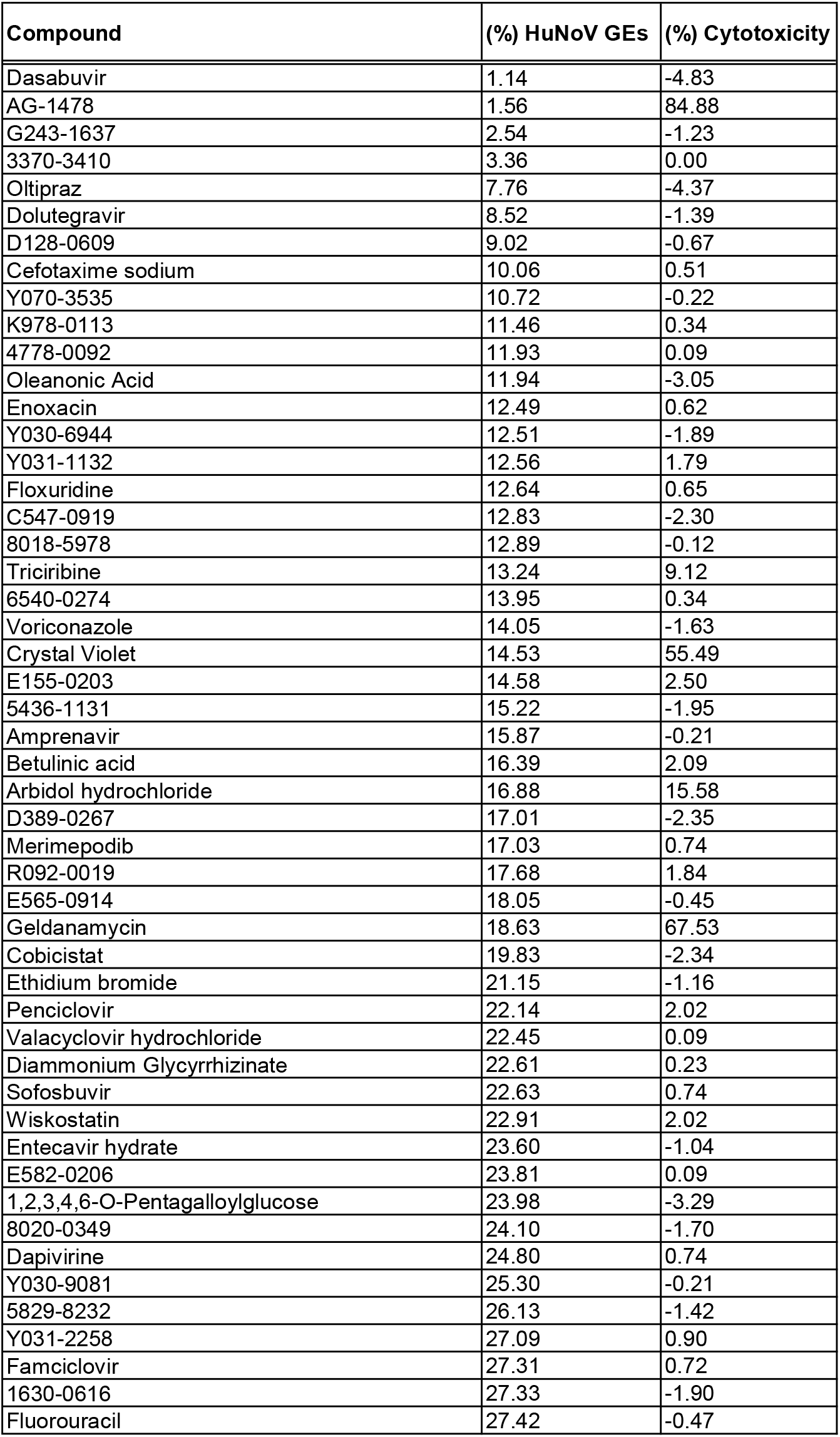
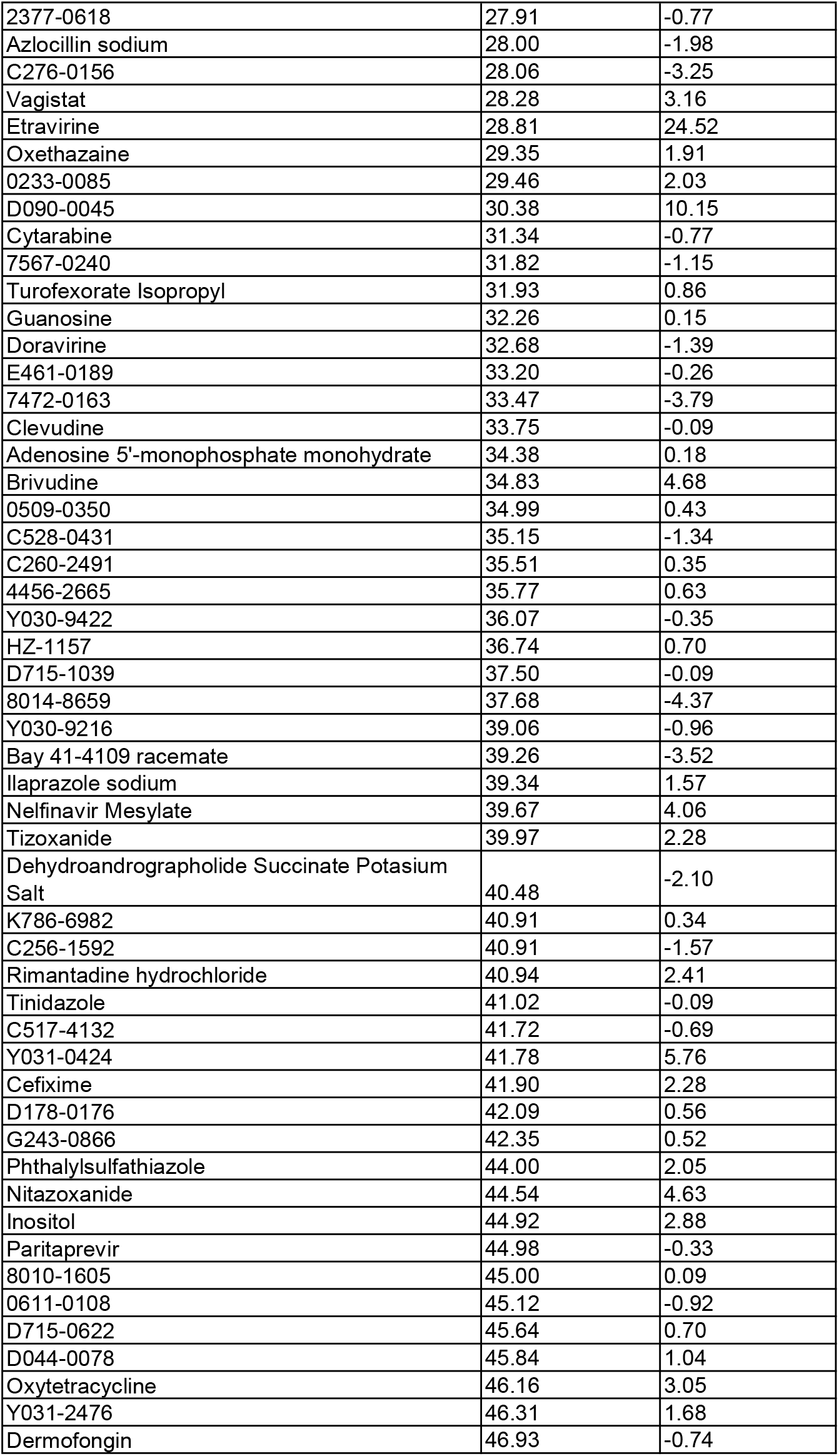
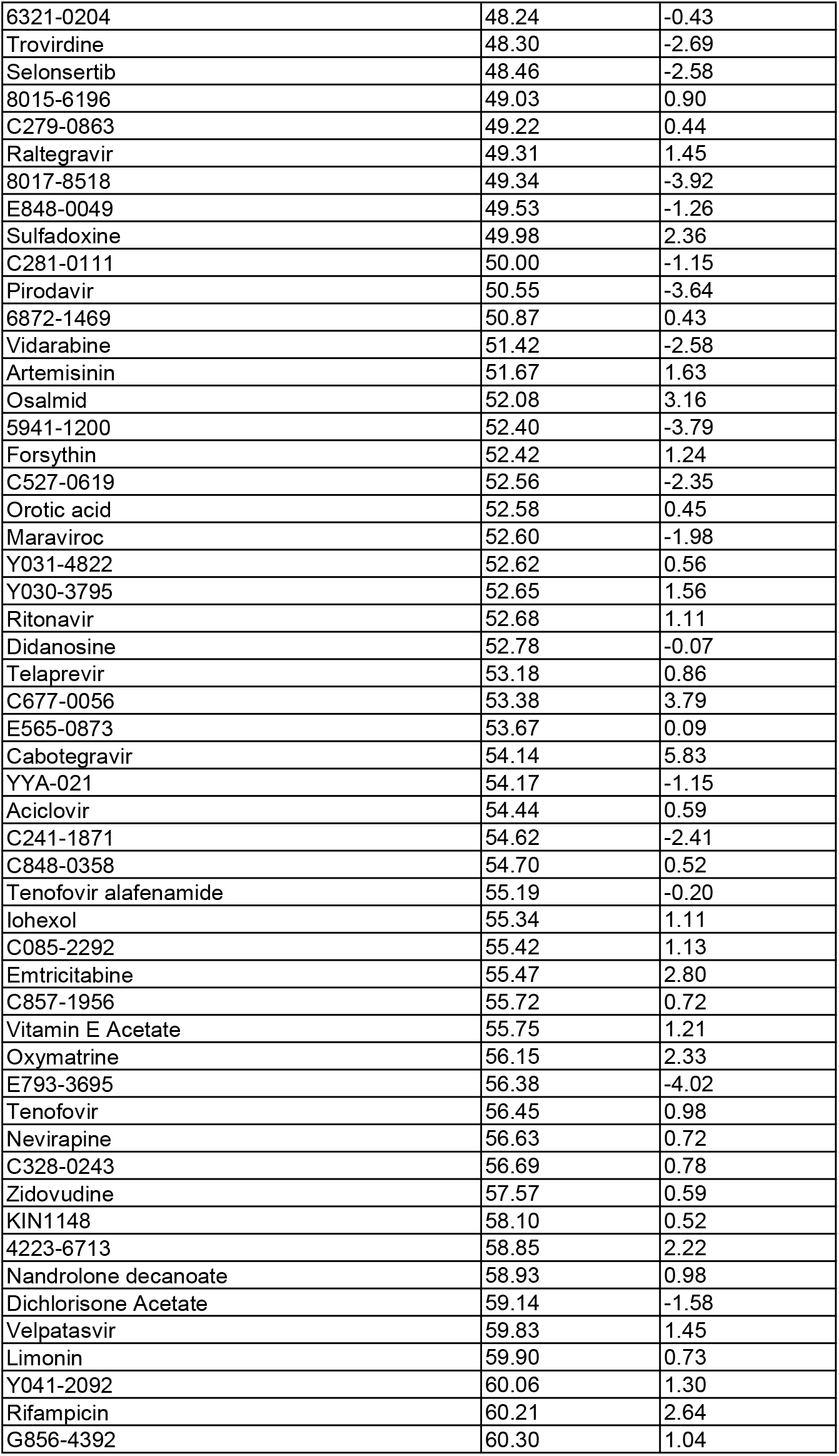
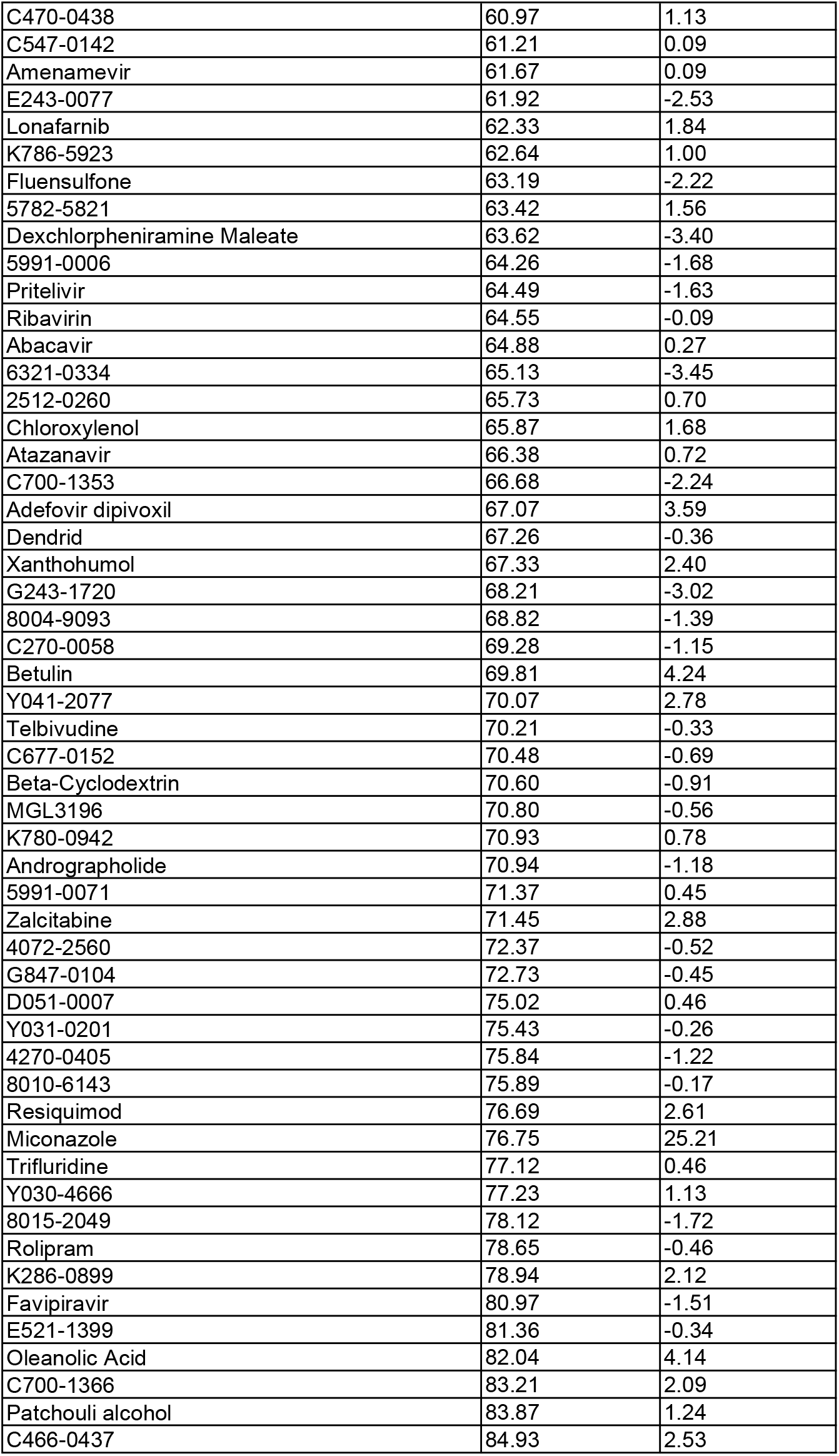
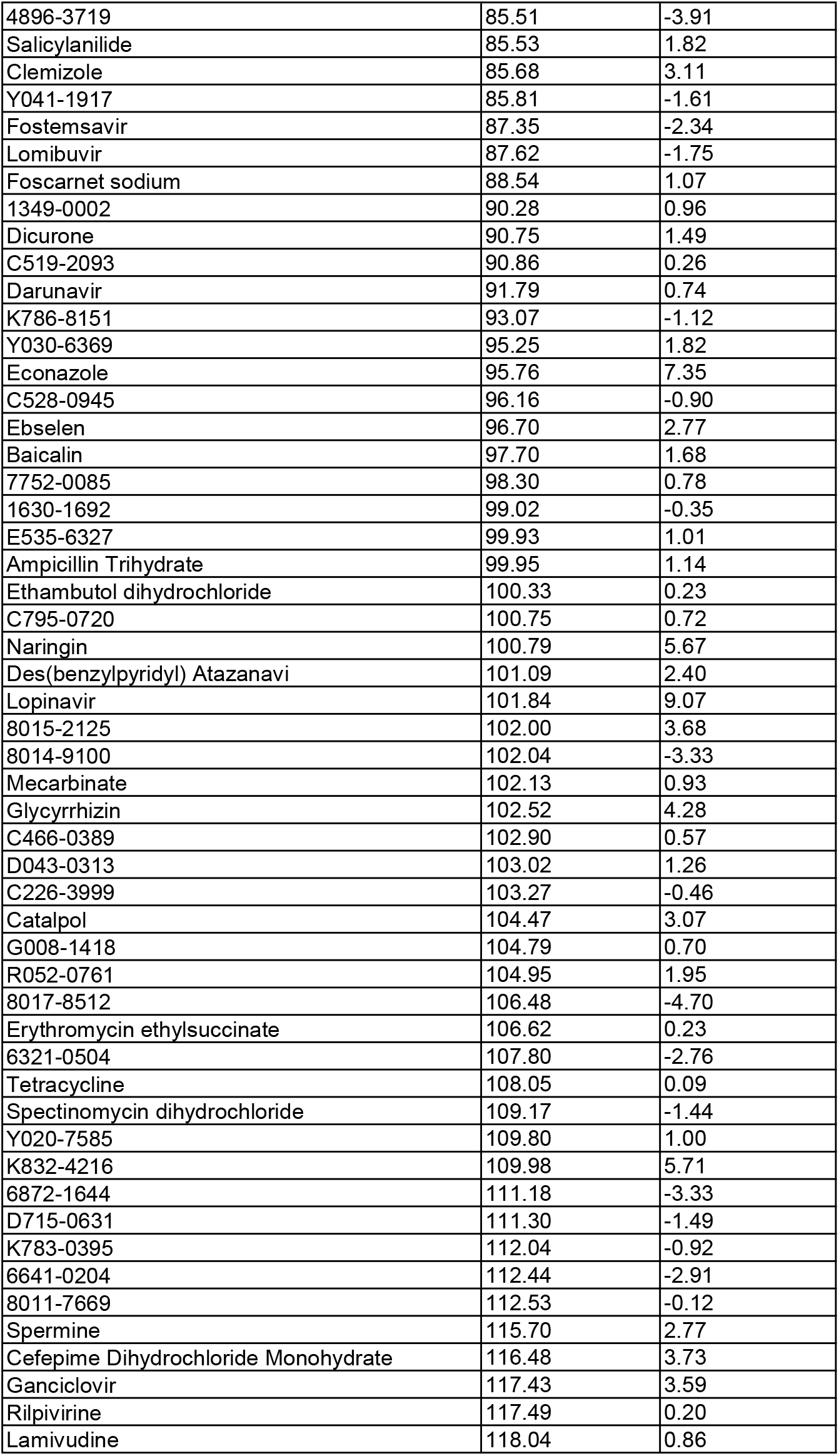
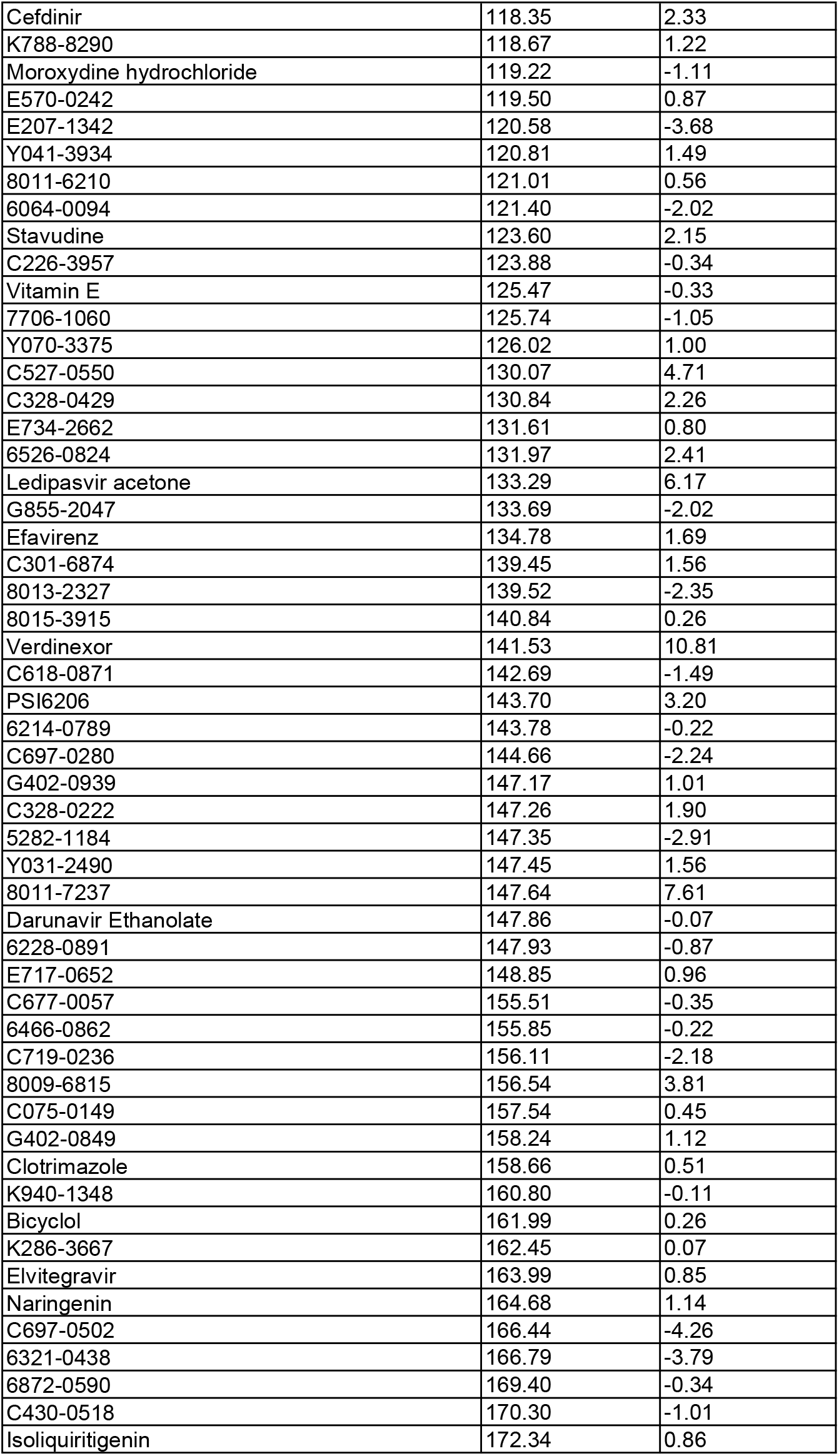
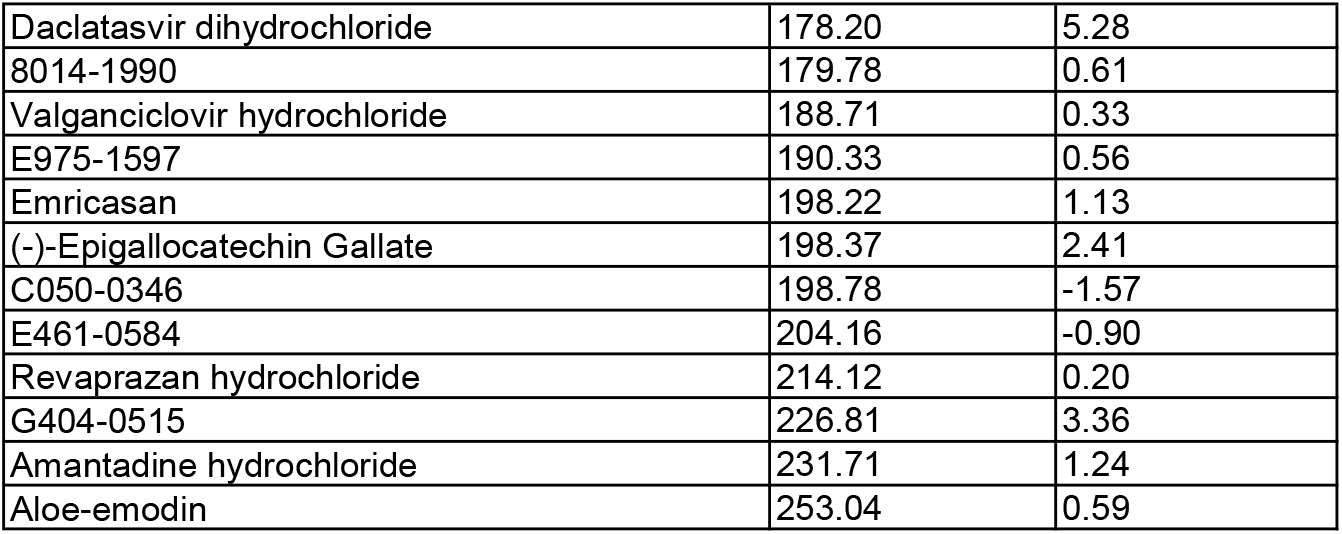
Compound screening results. The percentages of HuNoV GEs and cytotoxicity in the presence of the indicated compound (10 μM, n=1) were determined and normalized to those of DMSO control.

