## Supplemental material for "Dasabuvir inhibits human norovirus infection in human intestinal enteroids"

### Material and methods

#### Cells

Human intestinal enteroids (HIEs), J2 and J3 lines established from adult jejunal biopsy specimens (Ettayebi et al., 2016), were provided from Baylor College of Medicine under the Material Transfer Agreement. The study protocol was approved by the Review Board of National Institute of Infectious Diseases. Wnt3a-producing cells were kindly provided by Baylor College of Medicine. R-spondin- and Noggin-producing cells were kindly provided by Dr. Calvin Kuo (Palo Alto, CA, USA) and Dr. Gijs van den Brink (University of Amsterdam, Netherlands), respectively. HIEs were grown as multilobular, 3-dimensional (3D) cultures in Matrigel and were maintained in complete medium with growth factors [CMGF(+)] or IntestiCult Organoid Growth Medium (Human, Veritas) as previously described (Ettayebi et al., 2016; Ettayebi et al., 2021; Zou et al., 2019). A monkey kidney cell line MA104 was maintained in DMEM supplemented with 10% fetal bovine serum (FBS), 100 units/mL penicillin, and 100 μg/mL streptomycin. VeroE6/TMPRSS2 [JCRB1819, VeroE6 cell overexpressing the transmembrane protease, serine 2 (TMPRSS2)] (Matsuyama et al., 2020b) was purchased from JCRB Cell Bank (Osaka, Japan) and was maintained in DMEM supplemented with 10% FBS, 100 units/mL penicillin, 100 μg/mL streptomycin and 1 mg/mL G418 (Nacalai).

#### Viruses

Ten percent stool suspensions containing human norovirus (HuNoV) were prepared as described previously (Ettayebi et al., 2016) and were kept at -80 °C before use. Human A rotavirus (RVA) Wa strain was propagated in MA104 cells in the presence of trypsin.

Severe acute respiratory syndrome coronavirus 2 (SARS-CoV-2), 2019-nCoV/Japan/TY/WK-521/2020 strain (WK-521) was isolated previously (Matsuyama et al., 2020b) and was propagated in VeroE6/TMPRSS2 cells. Virus titer of the SARS-CoV-2 was determined by 50% tissue culture infectious dose ($TCID_{50}$) assay in VeroE6/TMPRSS2 cells (Matsuyama et al., 2020b).

**Compounds**

Anti-virus Compound Library (326 compounds, 10 mM solution in DMSO, L1700) was purchased from TargetMol. To validate antiviral effect, 3 individual dasabuvir were purchased from distinct companies (DSB, Cat. #T3489, TargetMol; Cat. #S5402, Selleck; Cat. #18482, Cayman Chemical). 2'-C-Methylcytidine (2-CMC, Cat. #22887) was purchased from Cayman Chemical. Remdesivir (RDV, Cat. #S8932) was purchased from Selleck.

**Infection of HIEs with viruses**

Virus infection in differentiated, two-dimensional (2D) HIE monolayers was performed following a previously described protocol with minor modification (Ettayebi et al., 2016; Ettayebi et al., 2021; Zou et al., 2019). Briefly, 3D HIEs were dissociated with TrypLE Express (Thermo Fisher) into single cells, after which they were seeded onto collagen IV-coated 96-well plates at the number of approximately ~$10^5$ cells/well in CMGF(+) or Intesticult media supplemented with ROCK inhibitor Y-27632 (10 μM, Sigma) for 2 days. After 2 days, the cells typically reach at ~100% confluency. The medium was then removed and the cells were maintained in the differentiation medium for another 2 days.

For HuNoV infection, the J2 and J3 HIE monolayers were inoculated with 10% stool filtrate containing 4.3 × $10^5$ genome equivalents (GEs) of GII.3 [GII.P21] (TCH04-577,(Ettayebi et al., 2016; Murakami et al., 2020) or GII.4 [GII.P16] HuNoV in the presence of 500 µM GCDCA, which is required for GII.3 HuNoV infection and promotes GII.4 infection (Ettayebi et al., 2016; Murakami et al., 2020). For RVA infection, the J2 HIE monolayers were inoculated with Wa strain at 2.48 × $10^8$ GEs/well that pre-treated with 10 µg/ml trypsin (Sigma) for 1 hr at 37 °C to activate VP4 spike protein (Crawford et al., 2001). For SARS-CoV-2 infection, the J2 monolayers were incubated with WK-521 at 8.1 × $10^8$ GEs/well. After 1 h incubation at 37 °C, the cells were washed twice with complete medium without growth factors [CMGF(-)] to remove unbound viruses. The cells were then incubated with differentiation medium in the presence (HuNoV) or absence (RVA and SARS-CoV-2) of GCDCA until 20-24 hrs post infection (hpi). The indicated compounds or DMSO were added to the medium throughout the infection process. The final concentration of DMSO was 0.5%. The cells and medium were then harvested and subjected to RNA extraction.

**RNA extraction and RT-qPCR**

Total RNA was extracted from the infected cells and/or media using Direct-zol RNA MiniPrep kit (Zymo Research), according to the manufacturer's instruction. RT-qPCR analysis to determine viral RNA GEs of HuNoV, RVA, and SARS-CoV-2 was performed using TaqMan Fast Virus 1-Step Master Mix (Thermo Fisher) and specific primer/probe sets as described previously (Ettayebi et al., 2016; Freeman et al., 2008; Matsuyama et al., 2020a).

**Cell viability assay**

Cytotoxicity or cell viability upon compound treatment was determined using Cytotoxicity LDH Assay Kit-WST (Dojindo) or CellTiter-Glo® Luminescent Cell Viability Assay (Promega), respectively, according to the manufacturer's instructions.

**Immunofluorescence assay**

Confluent VeroE6/TMPRSS2 cells cultured in 96 well plate (CellCarrier-96 Ultra, Perkin Elmer) were infected with SARS-CoV-2 at a multiplicity of infection (MOI) of 0.003 for 20 hrs at 37 °C. The infected cells were then fixed with 4% paraformaldehyde in D-PBS for 30 min and permeabilized with 0.2% Triton X-100 in D-PBS for 15 min. The cells were stained for SARS-CoV-2 Spike (S) protein using rabbit anti-SARS-CoV-2 Spike RBD monoclonal antibody (1:3,000, clone HL1003, GTX635792, GeneTex) followed by goat anti-rabbit IgG AlexaFluor 488 (1:1,000, Life technologies). Cell nuclei were stained with 1 μg/ml DAPI solution (Dojindo). The cells were then imaged using Operetta CLS High-Content Analysis System (Perkin Elmer) and the percentages of SARS-CoV-2 positive cells (infectivity) in each well were calculated by counting SARS-CoV-2 S and DAPI positive cells using Harmony software (Perkin Elmer).

**Data analysis and statistics**

All experiments except for compound screening (Figs. 1 and S1), were performed at least two times with more than two technical replicates and results are shown as the mean ± s.d. ($n \geq 4$). Statistical analysis was performed with ANOVA followed by Dunnett's multiple-comparison test or two-tailed Student *t* test using GraphPad Prism 9 software. *P* values of $< 0.05$ was considered statistically significant. Dose response curve was created by nonlinear regression model, and the 50% effective concentration ($EC_{50}$) and the cytotoxic concentration ($CC_{50}$) was calculated using GraphPad Prism 9 software.
